# Daisy-chain gene drives for the alteration of local populations

**DOI:** 10.1101/057307

**Authors:** Charleston Noble, John Min, Jason Olejarz, Joanna Buchthal, Alejandro Chavez, Andrea L. Smidler, Erika A. DeBenedictis, George M. Church, Martin A. Nowak, Kevin M. Esvelt

## Abstract

RNA-guided gene drive elements could address many ecological problems by altering the traits of wild organisms, but the likelihood of global spread tremendously complicates ethical development and use. Here we detail a localized form of CRISPR-based gene drive composed of genetic elements arranged in a daisy-chain such that each element drives the next. “Daisy drive” systems can duplicate any effect achievable using an equivalent global drive system, but their capacity to spread is limited by the successive loss of non-driving elements from the base of the chain. Releasing daisy drive organisms constituting a small fraction of the local wild population can drive a useful genetic element to local fixation for a wide range of fitness parameters without resulting in global spread. We additionally report numerous highly active guide RNA sequences sharing minimal homology that may enable evolutionary stable daisy drive as well as global CRISPR-based gene drive. Daisy drives could simplify decision-making and promote ethical use by enabling local communities to decide whether, when, and how to alter local ecosystems.

**Author’s Summary:** ‘Global’ gene drive systems based on CRISPR are likely to spread to every population of the target species, hampering safe and ethical use. ‘Daisy drive’ systems offer a way to alter the traits of only local populations in a temporary manner. Because they can exactly duplicate the activity of any global CRISPR-based drive at a local level, daisy drives may enable safe field trials and empower local communities to make decisions concerning their own shared environments.

For more details and an animation intended for a general audience, see the summary at <u>Sculpting Evolution</u>.

## Introduction

RNA-guided gene drive elements based on the CRISPR/Cas9 nuclease could be used to spread many types of genetic alterations through sexually reproducing species[1]. These elements function by “homing”, or the conversion of heterozygotes to homozygotes in the germline, which renders offspring more likely to inherit the gene drive element and the accompanying alteration than via Mendelian inheritance (Fig. 1a)[2]. To date, gene drive elements based on Cas9 have been demonstrated in yeast[3], fruit flies[4], and two species of mosquito[5][6]. Drive homing occurred at high efficiency (>90%) in all four species, strongly suggesting that refined versions may be capable of altering entire wild populations. Potential applications include eliminating vector-borne and parasitic diseases, promoting sustainable agriculture, and enabling ecological conservation by curtailing or removing invasive species.

The self-propagating nature of global gene drive systems renders the technology uniquely suited to addressing large-scale ecological problems, but tremendously complicates discussions of whether and how to proceed with any given intervention. Technologies capable of unilaterally altering the shared environment require broad public support. Hence, ethical gene drive research and development must be guided by the communities and nations that depend on the potentially affected ecosystems. Unfortunately, attaining this level of engagement and informed consent becomes progressively more challenging as the size of the affected region increases. Candidate applications that will affect multiple nations could be delayed indefinitely due to lack of consensus.

A method of confining gene drive systems to local populations would greatly simplify community-directed development and deployment while also enabling safe field testing. Existing theoretical strategies[7][8] can locally spread cargo genes nearly to fixation if sufficient organisms (>30% of the local population) are released. “Threshold-dependent” drive systems such as those employing underdominance[9] will spread to fixation in small and geographically isolated subpopulations if organisms exceeding the threshold for population takeover are released (typically ~50%). Toxin-based underdominance approaches are promising and have been demonstrated in fruit flies[10][11], but are more limited in their potential effects than are homing-based drive systems. All of these approaches involve releasing comparatively large numbers of organisms, which may not be politically, economically, or environmentally feasible.

A way to construct locally-confined RNA-guided drive systems could enable many potential applications for which neither global drive systems nor existing local drives are suitable. Here we describe ‘daisy drive’, a powerful form of local drive based on CRISPR-mediated homing in which the drive components are separated into an interdependent daisy-chain. We additionally report newly characterized guide RNA sequences required for evolutionary stability and safe use.

## Results

### Design and Modeling

**Figure 1:**
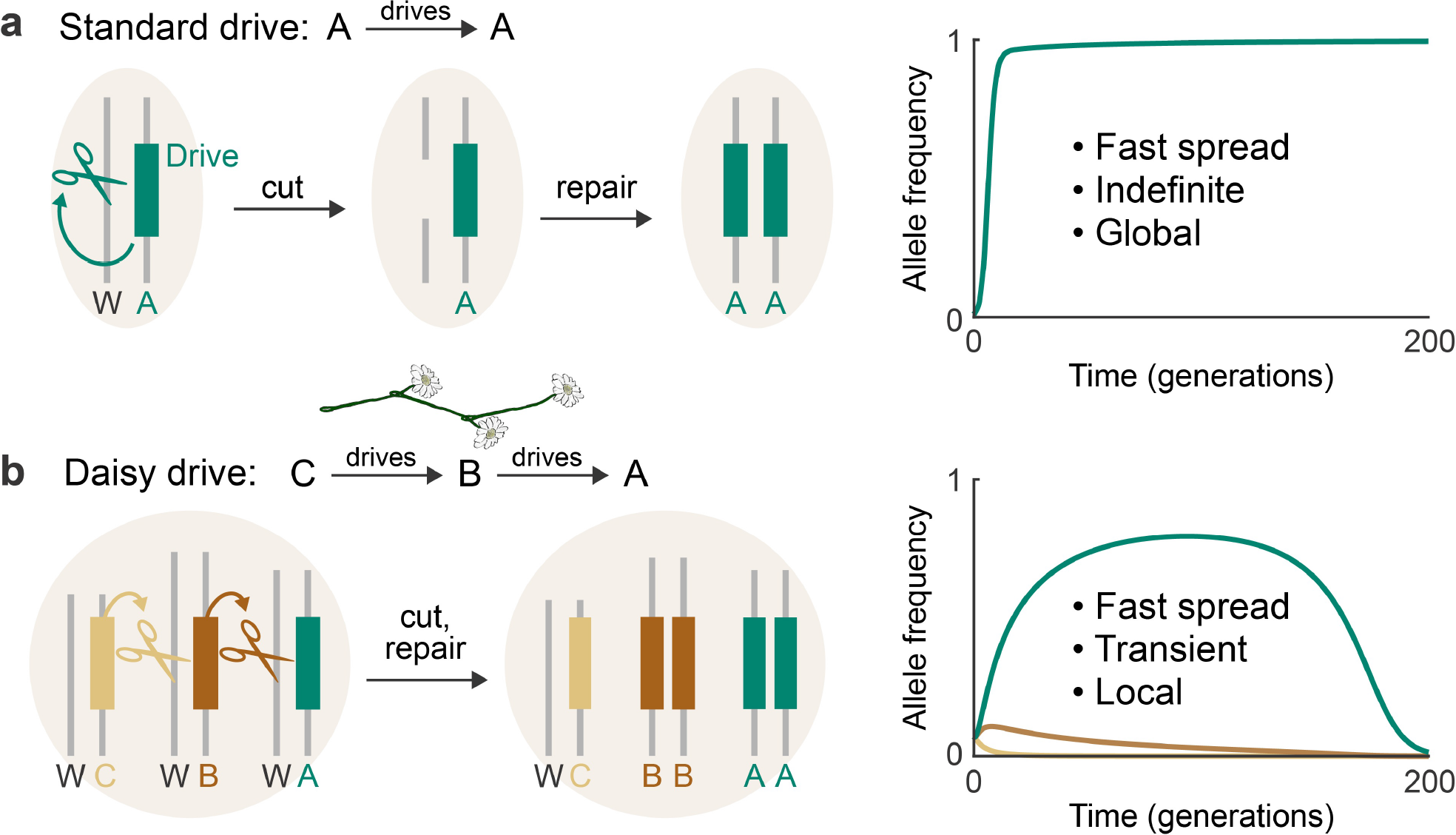
| a, Standard CRISPR gene drives distort inheritance in a self-sustaining manner by converting wild-type (W) alleles to drive alleles in heterozygous germline cells. b, A “daisy drive” system consists of a linear chain of serially dependent, unlinked drive elements; in this example, A, B, and C are on separate chromosomes. Elements at the base of the chain cannot drive and are successively lost over time via natural selection, limiting overall spread.

A daisy drive system consists of a linear series of genetic elements arranged such that each element drives the next in the chain (Fig. 1b). The top element, which carries the “payload”, is driven to higher and higher frequencies in the population by the elements below it in the chain. No element in the chain drives itself. The bottom element is lost from the population over time, causing the next element to cease driving and be lost in turn. This process continues up the chain until, eventually, the population returns to its wild-type state (Fig. 1b).

The simplest form of daisy drive—a two element chain—is obtained by separating CRISPR gene drive components such that the payload-carrying element, designated ‘A’, exhibits drive only in the presence of an unlinked, non-driving element, ‘B’ (Supplementary Fig. 1). These “split drives” have been described[1], demonstrated[3], and recommended[12] as a stringent laboratory confinement strategy. Because any accidental release would involve only a small number of organisms carrying the B element, the driving effect experienced by the A element—and thus its spread—would be negligible in a large population[3]. As long as the payload confers a fitness cost to the host organism, both elements will eventually disappear due to natural selection.

We hypothesized that the spread of the payload-carrying element, A, could be enhanced by adding more elements to the base of the daisy chain. To explore this idea, we formulated a deterministic model which considers the evolution of a large population of diploid organisms affected by a daisy drive system with elements spread acrossn loci (Supplementary Methods Section 1). At each locus there are two alleles, the wild-type (W) and the corresponding daisy drive element (D). To model the effects of drive in individuals, we assume that germline cells which are heterozygous at a locus convert to drive-homozygotes at that locus with probability *H* if the previous locus has at least one copy of a drive allele. In other words, individuals with genotype DW at locus *i*+1 and at least one copy of D at locus *i* produce gametes having the D allele at *i*+1 with probability (1+*H*)/2. We assume that standard Mendelian inheritance occurs in the absence of drive and that all loci are unlinked (e.g., on different chromosomes). We ignore the possible emergence of drive-resistant alleles because these can be prevented by ensuring that each element targets an essential gene with multiple guide RNAs and replaces it with a recoded version[1][13].

To model selection dynamics, we assumed that each construct confers a dominant fitness cost, q, on its host organism and that these costs are independent (Supplementary Fig. 2; SM Section 1.3). We assume that the target gene—a recoded copy of which is also contained in the corresponding drive element—is haploinsufficent. In this scenario, if a locus *i* contains a drive element and the next locus does not, then the drive cuts both wild-type alleles at that locus until both copies are disrupted, rendering resulting gametes nonviable (*c_i_* = 1). If the next locus instead contains two copies of the drive, then no cutting occurs (*c_i_* = 0). If there is exactly one drive allele at the next locus, then the wild-type allele is disrupted by cutting, rendering the organism nonviable unless a successful homing event occurs, in which case the drive is copied, a second copy of the target gene is created, and function is rescued. This occurs with probability *H*, so the associated cost is *c_i_* = 1-*H*.

Importantly, these costs are expected to be low because reported RNA-guided gene drives exhibit very high homing efficiencies: over 99% for each of the many drive systems tested in yeast[3], 95% for the fruit fly drive element[4], 99.8% for the drive element in *An. stephensi*[5], and 87.3% to 99.7% for the three drive systems in *An. gambiae*[6]. If the target gene is haploinsufficient for gametogenesis, the cost may even be zero (Supplementary Fig. 3). Finally, we assume that the payload element confers an additional dominant cost, *c_n_*, which is independent of this process. The total fitness of an individual is equal to *f* = (1-*c*_1_)(1-*c*_2_)…(1-*c_n_*).

An additional implicit assumption of our model for selection dynamics is that non-payload elements only confer costs via wild-type target gene disruption. We consider this reasonable because most elements in the daisy chain can consist of only guide RNAs, which should confer much lower costs than typical payloads[14][15]; moreover, potentially costly off-target cutting is minimal when using high-fidelity Cas9 variants[16] [17].

We studied a three-element daisy drive system (C→B→A) via numerical simulation (Fig. 2). We find that arbitrarily high frequencies of the payload element, A, can be achieved by varying the release frequency. However, the system displays high sensitivity to the homing rate and payload cost. In particular, large release sizes (>10% of the resident population) are required to drive costly payloads (>10%) if homing has efficiency on the lower end of observed drive systems (~90%).

**Figure 2:**
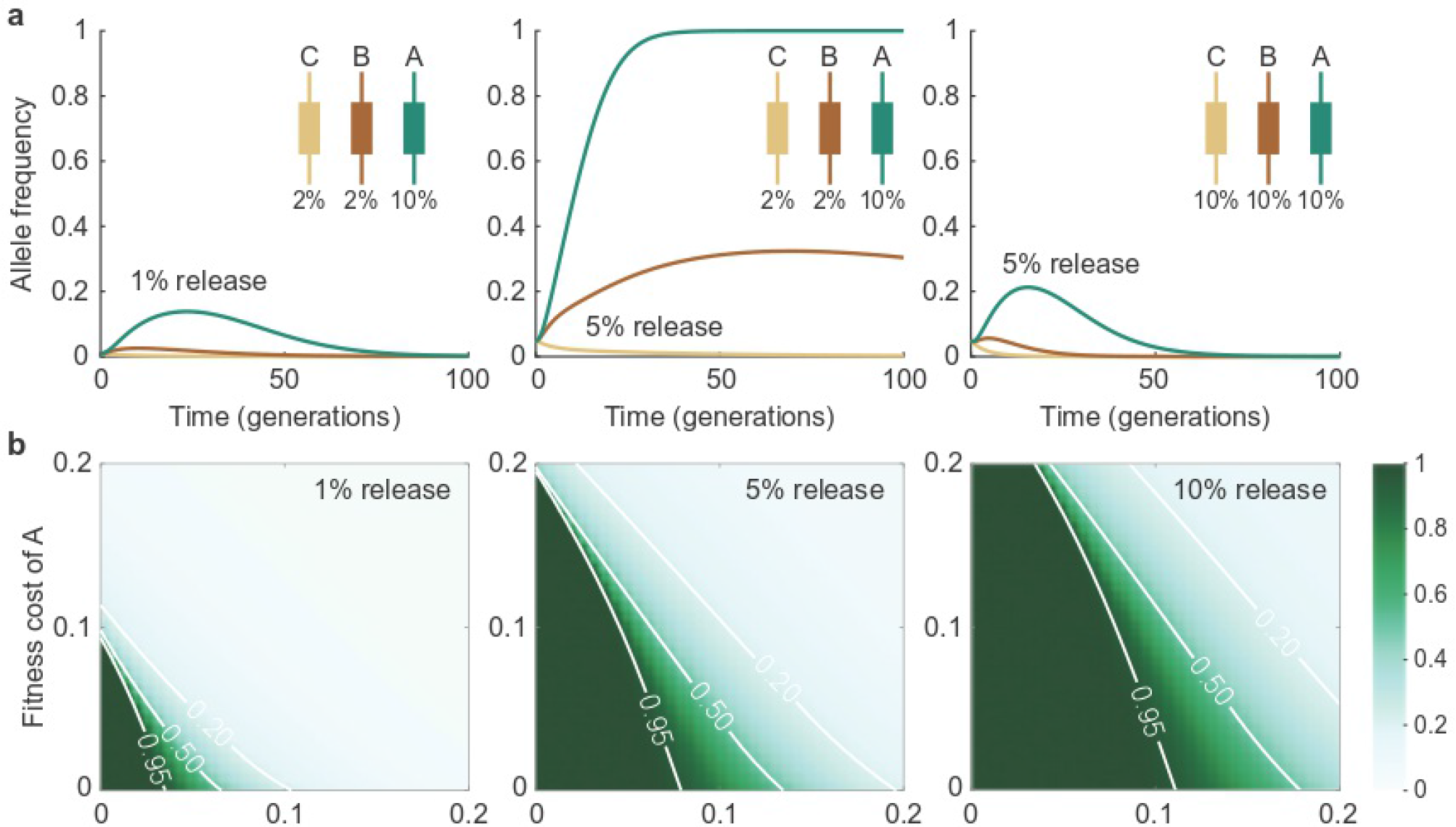
| Dynamics of C→B→A daisy drive systems. a, A highly efficient daisy drive (98% homing efficiency) with a 10% fitness cost for the payload element, seeded at 1%, exhibits limited spread (left). The same drive seeded at 5% rapidly spreads the payload to nearfixation (middle). Decreasing the homing efficiency to 90% would then require a larger release size (right). b, The maximum frequency achieved by C→B→A daisy drives as a function of the homing efficiency and the payload cost, for release sizes of 1% (left), 5% (middle), and 10% (right).

We next explored the effects of adding additional elements to the daisy drive system as a potential means of increasing their potency. We observe that longer chains lead to much stronger drive (Fig. 3). At a homing efficiency of 95% per daisy drive element, which is readily accessible to current drive systems, four-and five-element systems driving a payload with 10% cost could be released at frequencies as low as 5% and 3%, respectively, and still exceed 99% frequency in fewer than 20 generations. On a per-organism basis, these are over 100-fold more efficient than simply releasing organisms with the payload (Supplementary Fig. 4).

Adjusting the model to include repeated releases in every subsequent generation, we observed that daisy drives can readily alter local populations if repeatedly released in very small numbers, although the benefit of repeated release is lost when the initial release size becomes large (>10%) (Supplementary Fig. 5). This may be useful for applications that must affect large geographic regions over extended periods of time, as well as for local eradication campaigns[18].

**Figure 3:**
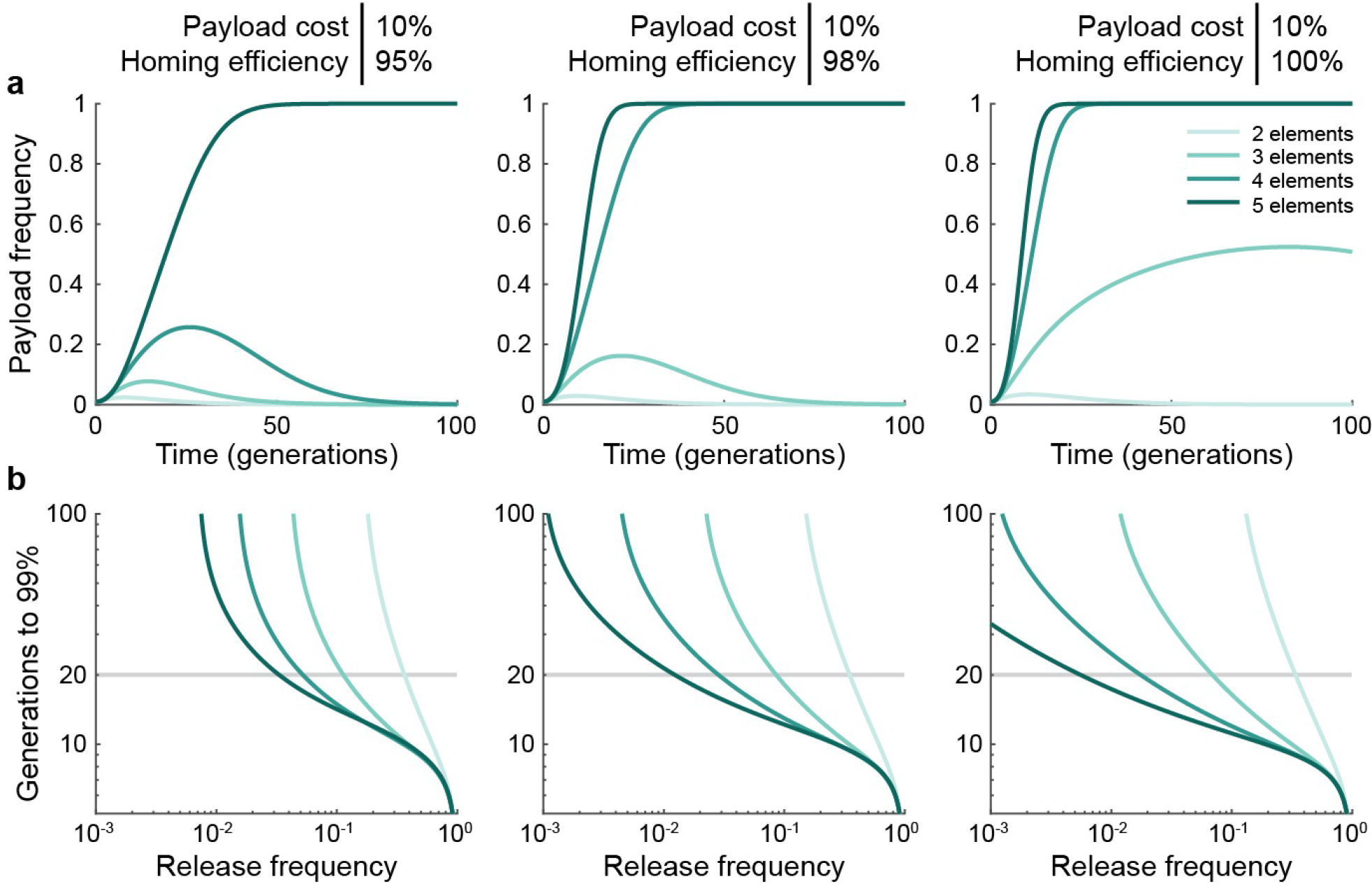
| The maximum frequency of the payload element (A), as well as its time to 99% frequency in a population, increases with the number of elements in the daisy chain. a, Example simulations assuming a 1% release of daisy drive organisms having a 10% payload fitness cost, and 95% (left), 98% (middle), or 100% (right) homing efficiencies. Darker shades indicate longer daisy chains (from 2 to 5 elements). b, Generations required for the payload element to attain 99% frequency.

## Evolutionary Stability and CRISPR Multiplexing

Despite these promising theoretical results, current technological limitations preclude the safe use of daisy drive elements. Specifically, any recombination event that moves one or more guide RNAs within an upstream element of the chain into any downstream element will convert a linear daisy drive chain into a self-sustaining gene drive ‘necklace’ anticipated to spread globally (Fig. 4a).

The only way to reliably prevent such events is to eliminate regions of homology between the elements. Promoter homology can be removed by using different U6, H1, or tRNA promoters to express the required guide RNAs[19][20][21]; if there are insufficient promoters then each can drive expression of multiple guide RNAs using tRNA processing[22][23] or by connecting a pair of sgRNAs by a short linker. However, each element must still encode multiple guide RNAs >80 base pairs in length in order to prevent the creation of drive-resistant alleles, precluding safe and stable daisy drive designs.

One alternative is to use a distinct orthogonal CRISPR system for every daisy element[24] (Supplementary Fig. 6). Unfortunately, enhanced-specificity variants are only available for the *S. pyogenes* Cas9, it is more difficult to find multiple promoters suitable for Cas9 expression than for guide RNA expression, and the fitness cost is likely higher than an equivalent guide RNA element. We accordingly sought to identify highly active guide RNA sequences with minimal homology to one another that could enable safe daisy drive using only a single CRISPR nuclease.

We compared known tracrRNA, crRNA, and alternative sgRNA sequences for CRISPR systems related to that of *S. pyogenes* to identify bases tolerant of variation within the sequence of the most commonly used sgRNA (Fig. 4b–c). We then created dozens of sgRNA variants designed to be as divergent from one another as possible. Assaying these using a sensitive tdTomato-based transcriptional activation reporter identified 15 different sgRNAs with activities comparable to the standard version (Fig. 4d). Activity increased with the length of the first stem in agreement with other reports (Supplementary Figs. 7–8) [25]. This set of minimally homologous sgRNAs can be used to construct stable daisy drive systems of up to 5 elements with 4 sgRNAs per driving element, and will also facilitate multiplexed Cas9 targeting in the laboratory by permitting the commercial synthesis of DNA fragments encoding many sequence-divergent guide RNAs. Future studies will need to examine the stability of the resulting daisy drive systems in large populations of animal models.

Importantly, our divergent guide RNAs will also enable global CRISPR gene drive elements to overcome the problem of instability caused by including multiple repetitive guide RNA sequences in the drive cassette[26], which in turn is required in order to overcome drive-resistant alleles[13]. Using non-repetitive guides may consequently allow stable and efficient global drive elements to affect every organism in the target population.

**Figure 4:**
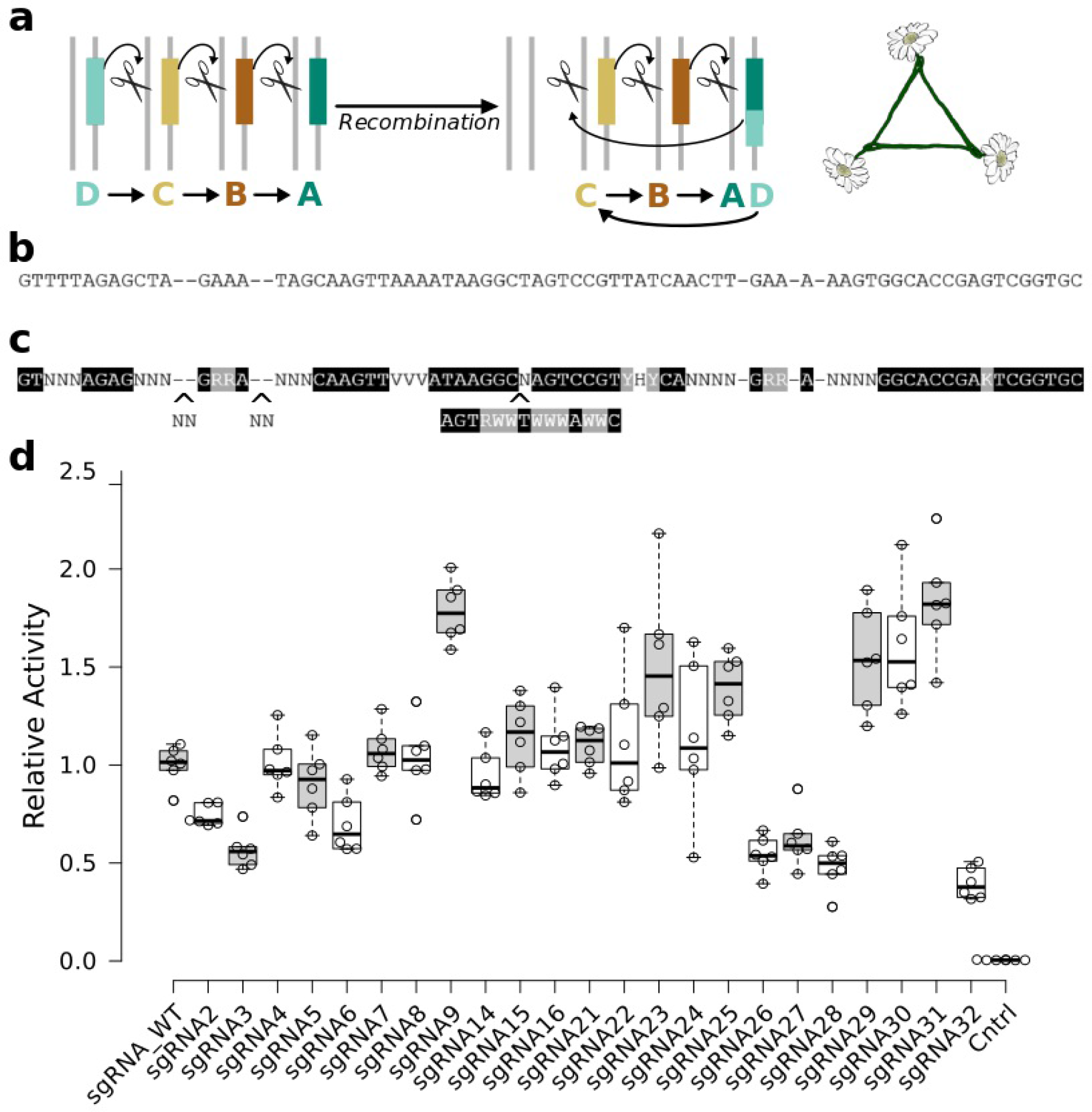
| a, Any recombination event that moves a guide RNA from one element to another could create a “daisy necklace” capable of self-sustaining global drive. b, Because promoters can be changed, repetition of the conserved guide RNA sequence is a key problem. c, Using existing data, we generated a template identifying candidate positions presumed tolerant of sequence changes. d, Relative activities of candidate guide RNAs generated from the template were assayed using a Cas9 transcriptional activator screen using a tdTomato reporter in human cells.

## Discussion

### Construction and Deployment

On a practical level, researchers need only construct one ‘generic’ daisy drive strain per species—the equivalent of a multistage rocket that could be loaded with any desired payload. This generic daisy drive system, which would harbor the Cas9 gene in the B position but lack any A elements, could be used in three different ways.

First, one or more elements carrying payloads could be added directly to the generic daisy drive strain. In addition to the payload, each such A element must encode guide RNAs sufficient to drive itself in the presence of Cas9. These daisy-drive organisms would then be mass-produced and released in a single-strain, single-stage approach.

Second, the generic daisy drive strain itself could be released in the target region to spread the Cas9 gene and accompanied by one or more strains carrying payload elements. Matings in the wild would combine the elements to generate the desired effect. This is a multi-strain, single-stage approach.

Third, the generic daisy drive strain could be released and the spread of the Cas9 gene monitored in order to identify the exact region that would be affected. Optionally, spread within this region could be adjusted by releasing wild-type organisms. Once acceptably distributed, a subsequent release of strains carrying payload elements would initiate the desired effect.

### Field Trials and Safeguards

Some ecological problems are so widely distributed geographically that addressing them may require global gene drive systems. However, global drive systems cannot be tested in field trials without a substantial risk of eventual worldwide spread[27]. Daisy drive systems, which are capable of mimicking the molecular effects of any given global drive on a local level, may offer a potential solution.

Similarly, scientists currently have few attractive options for controlling unauthorized or accidentally-released global drive systems. While it is possible to overwrite genome-level alterations and undo phenotypic changes using immunizing reversal drives[1], these countermeasures must necessarily spread to the entire population in order to immunize them against the unwanted drive system; strategies based on pure reversal drives[3] or variations such as gene drive ‘brakes’[28] will only slow it down. In contrast, daisy drive systems may be powerful enough to eliminate all copies of an unwanted global drive system via local immunizing reversal or population suppression before disappearing themselves.

Lastly, daisy drive systems could permit controlled and persistent population suppression by linking a sex-specific effect to a genetic locus unique to the other sex. For example, female fertility genes such as those recently identified in malarial mosquitoes[6] could be targeted by a genetic load daisy drive whose basal element is located on the Y chromosome or an equivalent male-specific locus (Supplementary Fig. 9). These males would suffer no fitness costs due to suppression relative to competing wild-type males. If female fertility gene disruption occurred early in development rather than in the germline, the same system could produce a male-linked dominant sterile-daughter effect that would be less powerful but more readily modulated (Supplementary Fig. 10).

By enabling scientists to reversibly control local population abundance, daisy drives could become a valuable tool for the study of ecological interactions and the likely consequences of releasing global RNA-guided suppression drives.

## Conclusion

RNA-guided gene drives based on CRISPR/Cas9 have generated considerable excitement as a potential means of addressing otherwise intractable ecological problems. While experiments have raced ahead at a breathtaking pace, the likelihood of global spread once released into the wild may prove a formidable barrier to deployment due to the need for international public support, field trials, and subsequent regulatory approval. These ethical and diplomatic complications are most acute for drive systems aiming to solve the most urgent humanitarian problems, including malaria, schistosomiasis, dengue, Zika, and other vector-borne and parasitic diseases. Lack of international consensus could delay approval by years or even decades.

Similarly, the potential for global RNA-guided drive systems to be released accidentally or deployed unilaterally has led to many calls for caution and expressions of alarm, not least from scientists in the vanguard of the field[1] [29][12]. Any such event could have potentially devastating consequences for public trust and support for future interventions.

In contrast, daisy drive systems might be safely developed in the laboratory, assessed in the field, and deployed to accomplish transient alterations that do not impact other nations or jurisdictions. By using molecular constraints to limit generational and geographic spread in a tunable manner, daisy drives could expand the scope of ecological engineering by enabling local communities to make decisions concerning their own local environments.

## Acknowledgments

We thank M. Tuttle for performing preliminary guide RNA activity assays, F. Gould, A. Lloyd, and L. Alphey for helpful discussions, and L. Alphey for critical reading of the manuscript.

## Author contributions

K.M.E. conceived the study, J.M. and K.M.E. ran preliminary simulations with advice from A.L.S.; C.N., J.O., M.A.N. created the evolutionary dynamics model, J.M. and K.M.E. designed the guide RNA template and candidate sequences, J.B. and A.C. designed and performed guide RNA experiments with advice from K.M.E., E.D. created the interactive version of the model, and C.N., J.M., and K.M.E. wrote the manuscript with contributions from all other authors.

## Methods

### Guide RNA Design

We examined existing data on guide RNA variants and corresponding activities as well as the crystal structure of *S. pyogenes* Cas9 in complex with sgRNA to identify bases that would likely tolerate mutation. Using this information, we constructed a set of 20 sgRNAs and assayed activity (see below) using only two replicates to identify sequence changes that were harmful to activity. These experiments suggested that the large insertion found in sgRNAs from closely related bacteria was well-tolerated in only one case. It was consequently removed and additional sgRNAs designed. All candidates were then assayed to identify those with sufficiently high activity. Future experiments requiring additional highly divergent sgRNAs, such as daisy suppression drives in which the A element encodes many guide RNAs that disrupt multiple recessive fertility genes at multiple sites, will require a more comprehensive library-based approach to activity profiling.

### Measuring Guide RNA Activity

HEK293T cells were grown in Dulbecco’s Modified Eagle Medium (Life Technologies) fortified with 10% FBS (Life Technologies) and Penicillin/Streptomycin (Life Technologies). Cells were incubated at a constant temperature of 37°C with 5% CO2. In preparation for transfection, cells were split into 24-well plates, divided into approximately 50,000 cells per well. Cells were transfected using 2ul of Lipofectamine 2000 (Life Technologies) with 200ng of dCas9 activator plasmid, 25ng of guide RNA plasmid, 60ng of reporter plasmid and 25ng of EBFP2 expressing plasmid.

Fluorescent transcriptional activation reporter assays were performed using a modified version of addgene plasmid #47320, a reporter expressing a tdTomato fluorescent protein adapted to contain an additional gRNA binding site 100bp upstream of the original site. gRNAs were co-transfected with reporter, dCas9-VPR, a tripartite transcriptional activator fused to the C-terminus of nuclease-null *Streptococcus pyogenes*Cas9, and an EBFP2 expressing control plasmid into HEK293T cells. 48 hours post-transfection, cells were analyzed by flow cytometry. In order to exclusively analyze transfected cells, cells with less than 10^ 3 EBFP2 expression were ignored. The preliminary screen of the initial 20 designs was performed with only two replicates to identify critical bases. Experiments evaluating the final set of sgRNA sequences were performed with six biological replicates.

## supplementary Methods

In these Supplementary Methods, we study the evolutionary dynamics of “daisy drive” genetic constructs. The aim of engineering such constructs is to genetically manipulate some—but not all—of a wild population.

### 1 Evolutionary dynamics of a daisy drive construct

We first describe a daisy drive system consisting of only two elements. This simple case demonstrates the principle behind daisy drive engineering. We then describe a daisy drive system with an arbitrary number of elements.

#### 1.1 Model for the evolutionary dynamics of a 2-element daisy drive

We consider a wild population of diploid organisms and focus on two loci, “ 1” and “2”. The wild-type alleles at the two loci are 1_*W*_ and 2_*W*_, and we denote by 1_*ww*_2_*ww*_ the genotype of an individual which is homozygous for both. Using CRISPR genome editing technology, one can engineer what we refer to as “daisy” alleles at both loci (1_*D*_ and 2_*D*_). They function as follows. The 1_*D*_ allele effects cutting of the 2*w* allele in an individual’s germline. After cutting, if the individual additionally has a 2_*D*_ allele (i.e., the individual is 2_*WD*_), then the individual is converted from 2_*WD*_ to 2_*DD*_ with some probability. Otherwise the individual remains 2_WD_. This results in super-Mendelian inheritance of the 2_*D*_ allele in a 1_*D*_-mediated fashion. Importantly, the 1_*D*_ allele undergoes standard inheritance and does not facilitate its own spread similarly. We assume that the two loci are independent and that a single copy of 1_*D*_ produces cutting.

To see how the daisy drive works, consider Table 1, which is understood as follows:

Gametes of haplotype 1_*w*_2_*W*_ are produced in the following ways:

- 1_*ww*_2_*WW*_ individuals produce only 1_w_2_W_ gametes.
- 1_*ww*_2_*WD*_ individuals produce gametes with a wild-type allele at the second locus with probability 1/2. There is a fitness effect, *F*, due to the payload of the drive allele at the second locus. So 1_ww_2_*WD*_ individuals produce 1_*w*_2_*W*_ gametes at relative rate *F*/2.
- 1_*WD*_2_*WD*_ individuals produce gametes with a wild-type allele at the first locus with probability 1/2. The action of the drive allele at the first locus is to bias the inheritance of the drive allele at the second locus, quantified by a factor *f_w_* regarding the transmission of the wild-type allele at the second locus. There is a fitness effect, *F*, due to the payload of the drive allele at the second locus. So 1_*WD*_2_*WD*_ individuals produce 1_*W*_2_*W*_ gametes at relative rate f_W_*F*/2.

**Table 1:** Gamete production table showing the relative rates at which individuals of each genotype (rows) produce gametes of each haplotype (columns).

| Genotype | $1_W 2_W$ | $1_W 2_D$ | $1_D 2_W$ | $1_D 2_D$ |
| --- | --- | --- | --- | --- |
| $1_{WW} 2_{WW}$ | 1 | 0 | 0 | 0 |
| $1_{WW} 2_{WD}$ | $\frac{1}{2}F$ | $\frac{1}{2}F$ | 0 | 0 |
| $1_{WW} 2_{DD}$ | 0 | $F$ | 0 | 0 |
| $1_{WD} 2_{WW}$ | 0 | 0 | 0 | 0 |
| $1_{WD} 2_{WD}$ | $\frac{1}{2}f_W F$ | $\frac{1}{2}f_D F$ | $\frac{1}{2}f_W F$ | $\frac{1}{2}f_D F$ |
| $1_{WD} 2_{DD}$ | 0 | $\frac{1}{2}F$ | 0 | $\frac{1}{2}F$ |
| $1_{DD} 2_{WW}$ | 0 | 0 | 0 | 0 |
| $1_{DD} 2_{WD}$ | 0 | 0 | $f_W F$ | $f_D F$ |
| $1_{DD} 2_{DD}$ | 0 | 0 | 0 | $F$ |

Gametes of haplotype 1_*W*_2_*D*_ are produced in the following ways:

- 1_*WW*_2_*WD*_ individuals produce gametes with a wild-type allele at the first locus with probability 1 and gametes with a drive allele at the second locus with probability 1/2. There is a fitness effect, *F*, due to the payload of the drive allele at the second locus. So 1_*WW*_2_*WD*_ individuals produce 1_*W*_2_D_ gametes at relative rate *F*/2.
- 1_WW_2_DD_ individuals produce only 1_*W*_2_*D*_ gametes. There is a fitness effect, *F*, due to the payload of the drive allele at the second locus. So 1_*WW*_2_*DD*_ individuals produce 1_*W*_2_*D*_ gametes at relative rate *F*.
- 1_*WD*_2_*WD*_ individuals produce gametes with a wild-type allele at the first locus with probability 1/2. The action of the drive allele at the first locus is to bias the inheritance of the drive allele at the second locus, quantified by a factor *f_D_* regarding the transmission of the drive allele at the second locus. There is a fitness effect, *F*, due to the payload of the drive allele at the second locus. So 1_WD_2_*WD*_ individuals produce 1_*W*_2_*D*_ gametes at relative rate *f*_*D*_*F*/2.
- 1_*WD*_2_DD_ individuals produce gametes with a wild-type allele at the first locus with probability 1/2 and gametes with a drive allele at the second locus with probability 1. There is a fitness effect, *F*, due to the payload of the drive allele at the second locus. So 1_*WD*_2_*DD*_ individuals produce 1_*D*_2_*W*_ gametes at relative rate *F*/2.

Gametes of haplotype 1_*D*_2_*W*_ are produced in the following ways:

- 1_*DD*_2_*WD*_ individuals produce gametes with a drive allele at the first locus with probability 1. The action of the drive allele at the first locus is to bias the inheritance of the drive allele at the second locus, quantified by a factor *f_W_* regarding the transmission of the wild-type allele at the second locus. There is a fitness effect, *F*, due to the payload of the drive allele at the second locus. So 1_*DD*_2_*WD*_ individuals produce 1_*D*_2_*W*_ gametes at relative rate *f_W_F*/2.
- 1_WD_2_WD_ individuals produce gametes with a drive allele at the first locus with probability 1/2. The action of the drive allele at the first locus is to bias the inheritance of the drive allele at the second locus, quantified by a factor *f_W_* regarding the transmission of the wild-type allele at the second locus. There is a fitness effect, *F*, due to the payload of the drive allele at the second locus. So l_*WD*_2_*DD*_ individuals produce l_*D*_2_*W*_ gametes at relative rate *fwF*/2.

Gametes of haplotype 1_*D*_2_*D*_ are produced in the following ways:

- 1_*WD*_2_WD_ individuals produce gametes with a drive allele at the first locus with probability 1/2. The action of the drive allele at the first locus is to bias the inheritance of the drive allele at the second locus, quantified by a factor *f_D_* regarding the transmission of the drive allele at the second locus. There is a fitness effect, *F*, due to the payload of the drive allele at the second locus. So 1_*WD*_2_*WD*_ individuals produce 1_*D*_2_*D*_ gametes at relative rate *f_D_F*/2.
- 1_*WD*_2_DD_ individuals produce gametes with a drive allele at the first locus with probability 1/2 and gametes with a drive allele at the second locus with probability 1. There is a fitness effect, *F*, due to the payload of the drive allele at the second locus. So 1_*WD*_2_DD_ individuals produce 1_*D*_2_*D*_ gametes at relative rate *F*/2.
- 1_*DD*_2_*WD*_ individuals produce gametes with a drive allele at the first locus with probability 1. The action of the drive allele at the first locus is to bias the inheritance of the drive allele at the second locus, quantified by a factor f_*D*_ regarding the transmission of the drive allele at the second locus. There is a fitness effect, *F*, due to the payload of the drive allele at the second locus. So 1_*DD*_2_*WD*_ individuals produce 1_*D*_2_*D*_ gametes at relative rate *f_D_F*.
- 1_*DD*_2_*DD*_ individuals produce only 1_*D*_2_*D*_ gametes. There is a fitness effect, *F*, due to the payload of the drive allele at the second locus. So 1_*DD*_2_*DD*_ individuals produce 1_*D*_2_*D*_ gametes at relative rate *F*.

Using these rules, we can formally express the rates at which the four types of gametes are produced in the population. We denote by *g*(*z*) the rate (with implicit time-dependence) at which gametes with haplotype *z* are produced by individuals in the population.

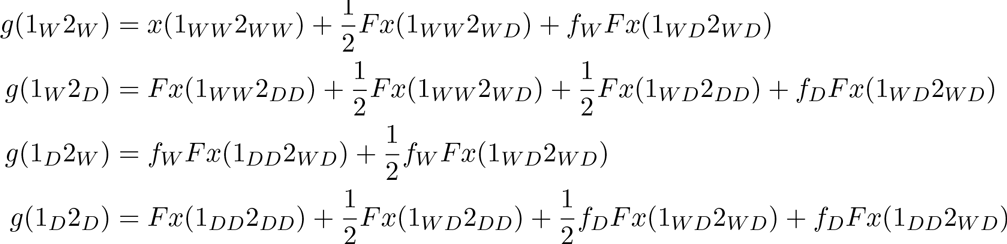

Here we have used the following notation: *x*(*z*) is the frequency of individuals with genotype *z* and *f*(*z*) is the fitness of individuals with genotype *z*.

The selection dynamics are then modeled by the following system of equations:

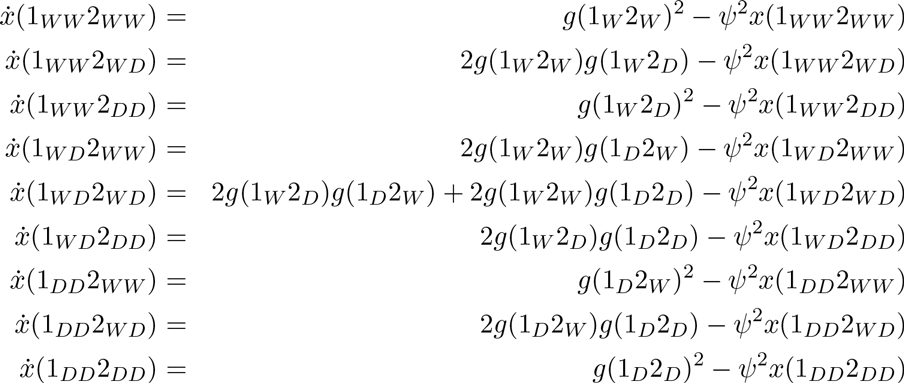

Here, an overdot denotes the time derivative, *d/dt.* Throughout this SI, we omit explicitly writing the time dependence of our dynamical quantities. Note that this formulation assumes random mating, i.e., that two random gametes come together to form an individual. Also note that products *g*(*y*)*g*(*z*) represent the pairings of different gametes. At any given time, we require that the total number of individuals sums to one:

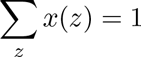

To enforce this density constraint, we set

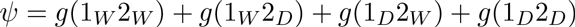

### 1.2 Model for the evolutionary dynamics of an *n*-element daisy drive

We can apply the same engineering to a daisy drive chain of arbitrary length *n*, where the drive allele at one locus induces cutting of the wild-type allele at the next locus in the sequence. To describe this mathematically, it is helpful to generalize our notation.

Consider a daisy drive construct with only two loci, as in the previous section. We use a “1” bit to denote a wild-type allele, and we use a “0” bit to denote a daisy drive allele. The two alleles at a particular locus are denoted in a vertical pair. Thus, we denote the nine possible genotypes as

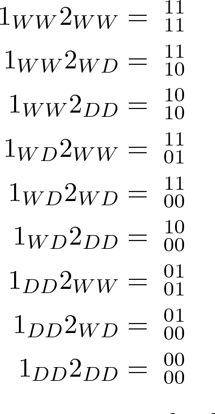

Notice that if an individual is heterozygous at a particular locus, then this notation allows for two ways of writing the alleles at that locus. For example, genotype 1 _*WD*_2_*WD*_ can be written in any one of four equivalent ways: 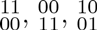, or 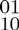.

When modeling daisy drives with a large number of loci, it is helpful to adopt shorthand notation. We use *p* and *q* to denote two binary strings of alleles. *p* and *q* need not be equal (i.e., the bits at corresponding positions need not all be equal). For example, genotype 1_*WW*_2_*WD*_ can be written as 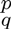, where *p* = 11 and *q* = 10. In this case, we denote the individual bits in the strings *p* and *q* by *p*_1_ = 1, *p*_2_ = 1, *q*_1_ = 1, and *q*_2_ = 0. Notice that *p* = 10 and *q* = 11 (i.e., *p*_1_ = 1, *p*_2_ = 0, *q*_1_ = 1, and *q*_2_ = 1) also describe the same genotype.

We denote by 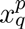 the frequency of individuals with genotype 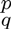. We denote by *g_q_* the rate at which gametes with haplotype *q* are produced. For an n-element daisy drive, *g_q_* is given by

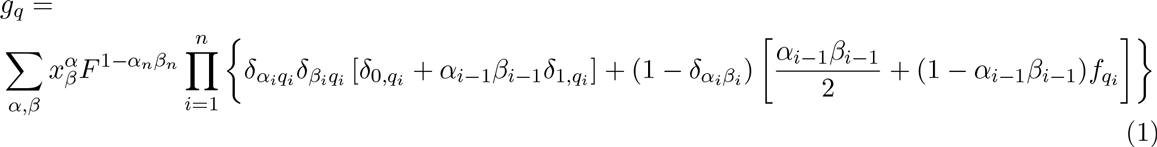

Here, we have defined *α1_0_* = *β*_0_ = 1. In the sum over α,β when enumerating genotypes, heterozygous loci (α_i_ ≠ β_i_are each counted once, so there is no double-counting. *g_q_* is linear in each 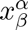, where all genotypes 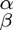 are summed over.

We understand the terms in the factors in brackets as follows. Consider just a single factor in brackets for a particular value of *i*.

- If *α_i_* = *β_i_* = *q_i_* = 0, then individuals of genotype 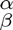 have two identical copies of allele 0 at the *i*^th^ locus, and those individials create only gametes with allele 0 at position *i*.
- If *α_i_* = *β_i_*= *q_i_* = 1 and *α_i-1_*, *β_i-1_* = 1, then individuals of genotype 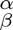 have two identical copies of allele 1 at the i^th^ locus and no copy of allele 0 at the (*i* − 1)^th^ locus, and those individials create only gametes with allele 1 at position *i*.
- If *a_i_* = β_i_ = *q*_i_, = 1 and *α_i-1_* β_i-1_ = 0, then individuals of genotype 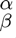 have two identical copies of allele 1 at the *i*^th^ locus and at least one copy of allele 0 at the (*i* − 1)^th^ locus. Since the daisy drive allele at the (*i* − 1)^th^ locus destroys both copies of the wild-type allele at the i^th^ locus, these individuals do not produce viable gametes. Hence, there is no corresponding term in Equation (1).
- If *α_i_* = *β_i_* and *α_i-1_*, *β_i-1_* = 1, then individuals of genotype 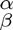 have a single copy of allele *q_i_* at the *i*^th^ locus, and without any action from the daisy drive, those individials create gametes with allele *q_i_* and allele (1 + (—1)^*qi*^)/2 at position *i* in equal proportion.
- If *α_i_* = *β_i_* and *α_i-1_*, *β_i-1_* = 0, then individuals of genotype 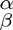 have a single copy of allele *q_i_* at the i^th^ locus, and the daisy drive allele at the (*i* − 1)^th^ locus biases the inheritance of allele *q_i_* at position *i*.

The prefactor *F^1-α_n_^β^n^*is the fitness cost associated with the payload. It appears if there is at least one copy of the daisy drive allele at the last position, *n*, in the daisy chain.

The selection dynamics for an *n*-element daisy drive are modeled by the following equations:

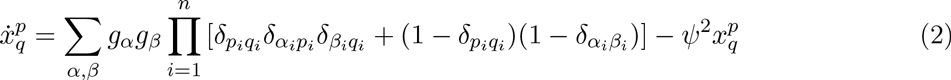

There is one such equation for each possible genotype 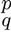.

We make sense of Equations (2) as follows. Each pair of gametes *g_α_* and g_β_ makes a new individual.

- If *p_i_* = *q_i_* = α_j_ = β_j_, then gametes of haplotypes a and pair to make only individials with genotype 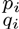 at locus *i*.
- If *p_j_* ≠ *q_j_* and *α_j_* ≠ β_*i*_ then gametes of haplotypes α and β pair to make only individials with genotype 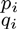 at locus *i*.

We impose the density constraint

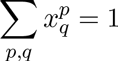

As already noted, in the sum over *p*, *q* when enumerating genotypes, heterozygous loci (*p_i_* ≠ *q_j_*) are each counted once, so there is no double-counting. We use the following identity

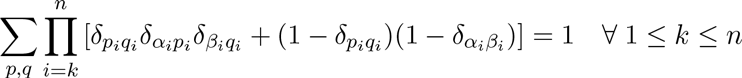

The form of *ψ* that enforces the density constraint is

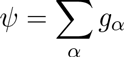

**Supplementary Figure 1.**
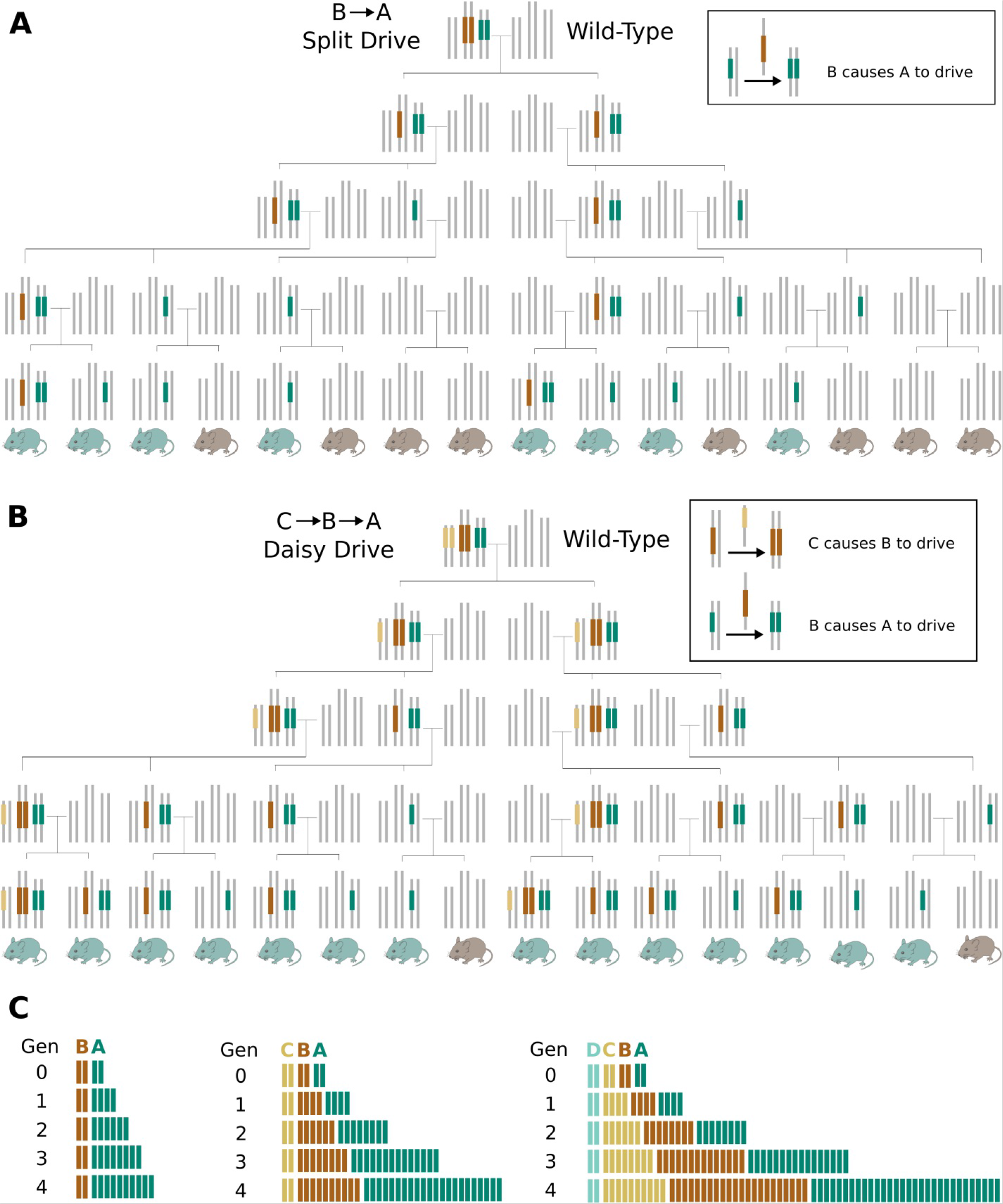
Family tree analysis of (A) B→A split drive and (B) C→B→A daisy drive demonstrates the power of including an additional daisy drive element in spreading the payload to more offspring in the F4 generation. (C) Graphical depiction of total alleles per generation for B->A through D->C→B→A daisy drives.

**Supplementary Figure 2.**
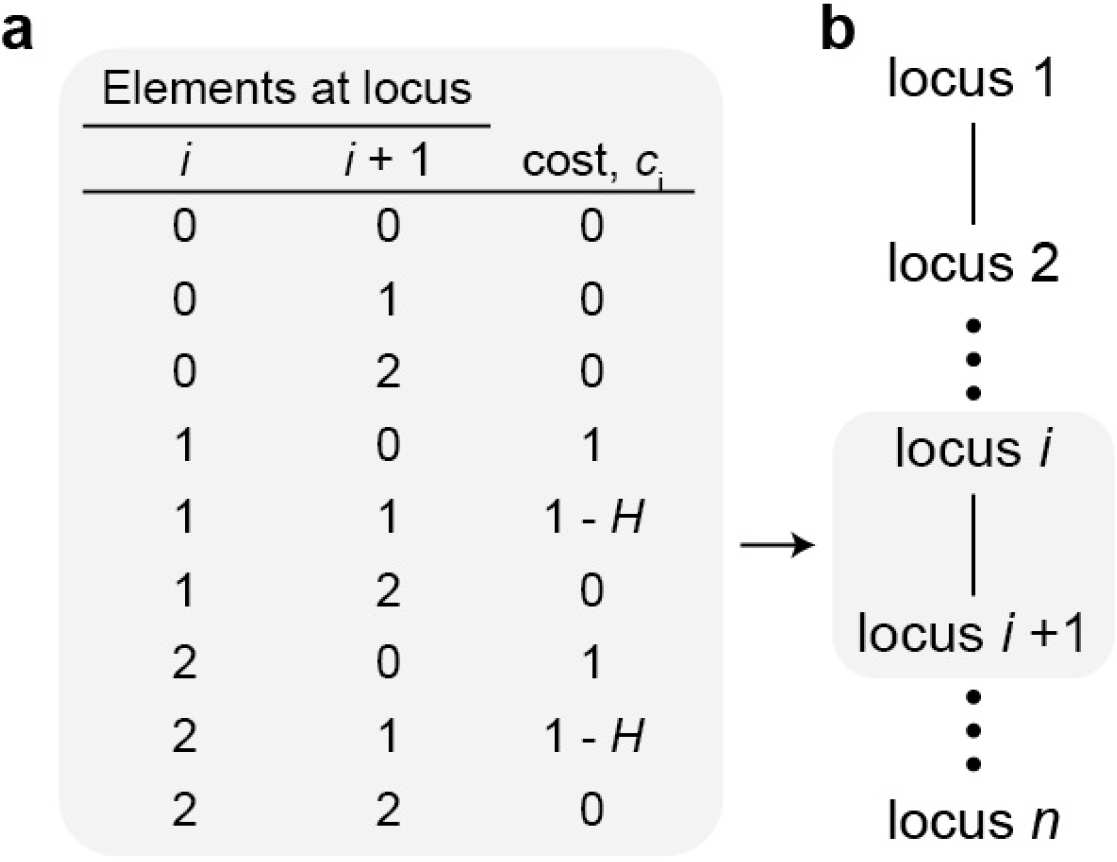
Mechanistic model for fitness parameters assumed in the daisy drive system. a, We assume that fitness costs primarily arise from cutting and misrepair events which result in disrupted haploinsufficient target genes. Such events occur only when there is a drive element at some locus (i) and a susceptible target at the next locus (i+1). If the next locus has two wild-type alleles—that is, no drive elements—then cutting and misrepair is lethal: the fitness cost associated with this pair of loci is thus ci = 1. If the next locus has one wild-type allele, then misrepair events occur precisely when homing does not succeed, and this happens with probability 1-H; thus the associated cost is ci = 1-H. If there is no drive element at the first position and/or no susceptible targets at the second position, then no cutting can occur and thus no cost is incurred. (b) We assume that the cost contributed by each link in the chain is independent. We calculate the costs ci for each pair of adjacent links, and the total fitness of the organism is then the product (1-c1)(1-c2)…(1-cn), where cn is the cost associated with the payload gene.

**Supplementary Figure 3.**
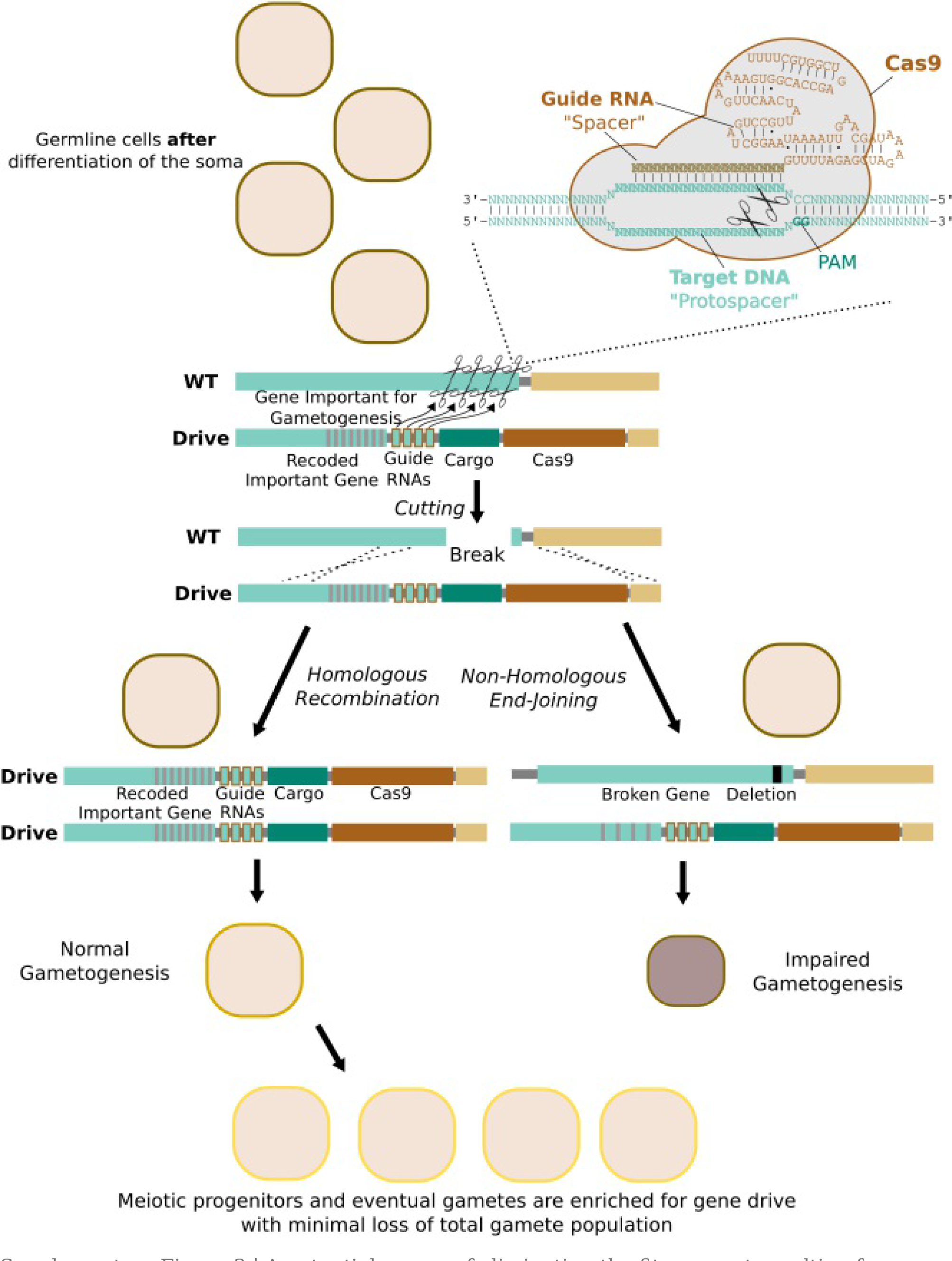
A potential means of eliminating the fitness cost resulting from incorrect repair might involves targeting a gene whose loss impairs gametogenesis, such as a ribosomal gene. Increased replication of correctly repaired cells carrying the drive system would theoretically result in a wild-type level of gametes, all of which carry the drive system.

**Supplementary Figure 4.**
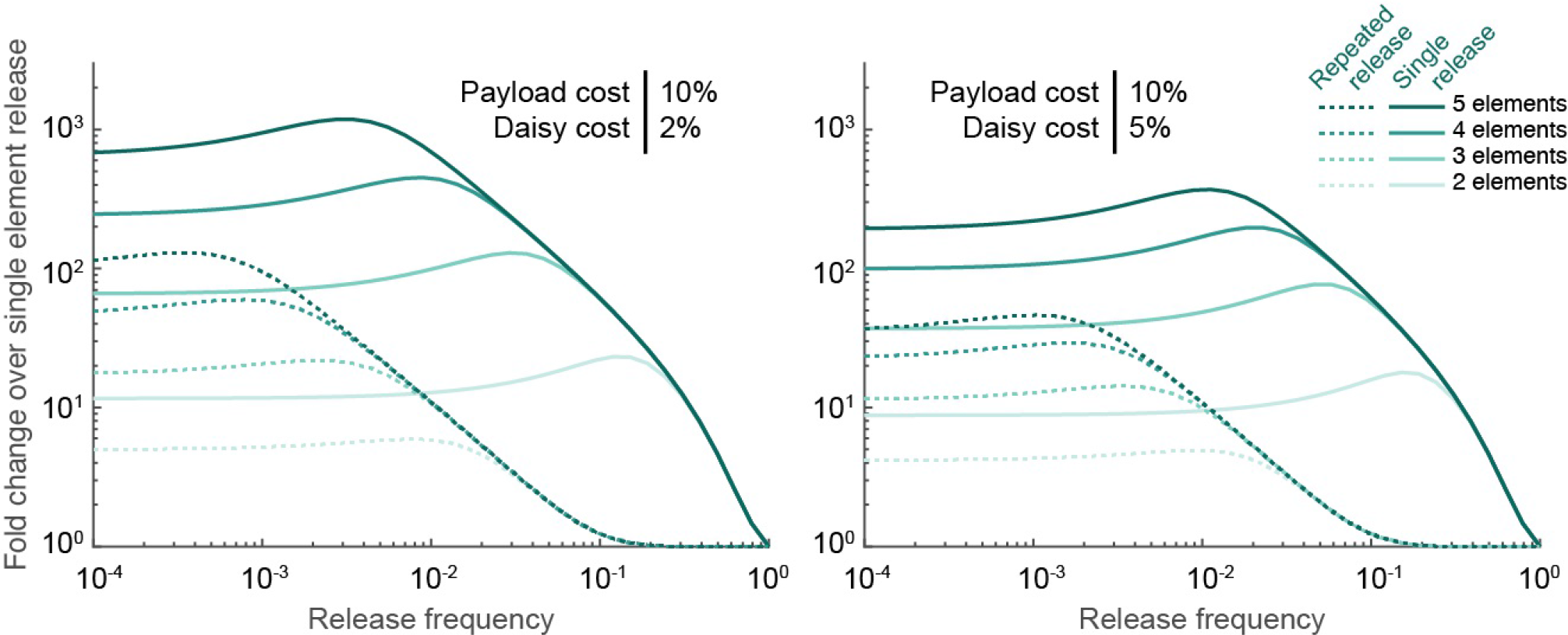
| The comparative efficacy of daisy drive systems can be assessed by comparing the payload frequency resulting from of releasing one daisy drive organism of different daisy-chain lengths after 2 0 generations relative to releasing organisms with only the payload.

**Supplementary Figure 5.**
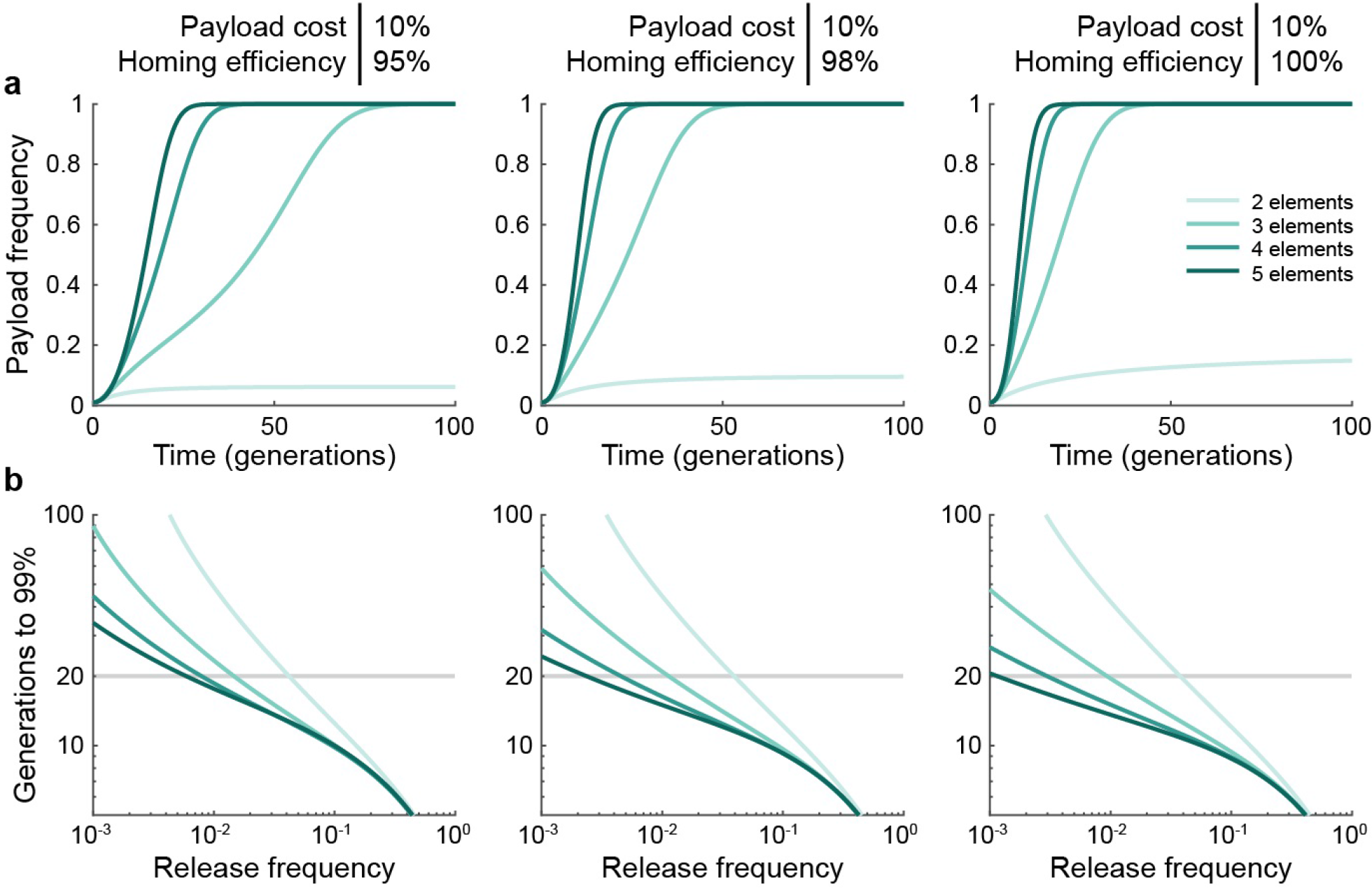
| Repeated seeding of engineered organisms improves daisy drive spread for low release frequencies. a, Example simulations assuming a 1% release of daisy drive organisms having a 10% payload fitness cost, and 95% (left), 98% (middle), or 100% (right) homing efficiencies. Darker shades indicate longer daisy chains (from 2 to 5 elements). b, Generations required for the payload element to attain 99% frequency. All simulations are identical to those in Fig. 3 of the main text, except here we assume that the initial release is repeated each generation.

**Supplementary Figure 6.**
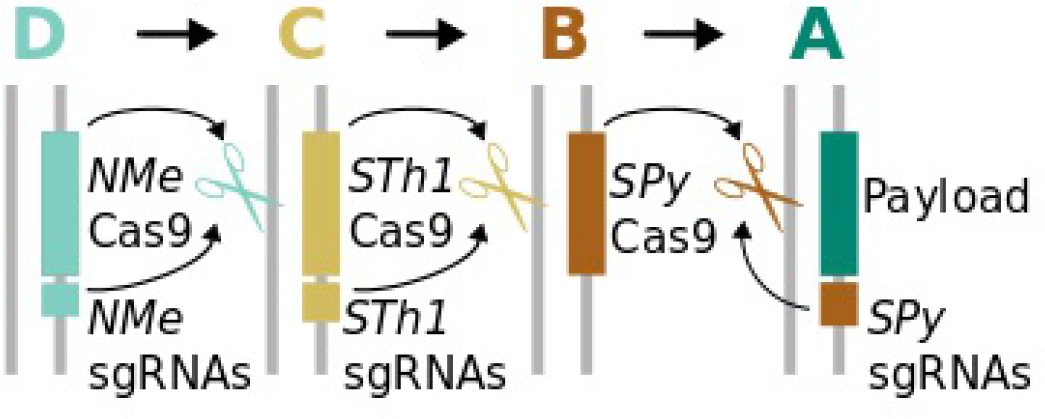
| Daisy drive systems can be constructed using orthogonal Cas9 elements. Such a drive system is resistant to conversion into a daisy necklace, which would require a recombination event that moved the entire Cas9 gene and associated guide RNAs into a subsequent locus in the daisy-chain. Ensuring that all the Cas9 proteins are expressed appropriately without re-using promoters and thereby creating homology between elements could be challenging.

**Supplementary Figure 7.**
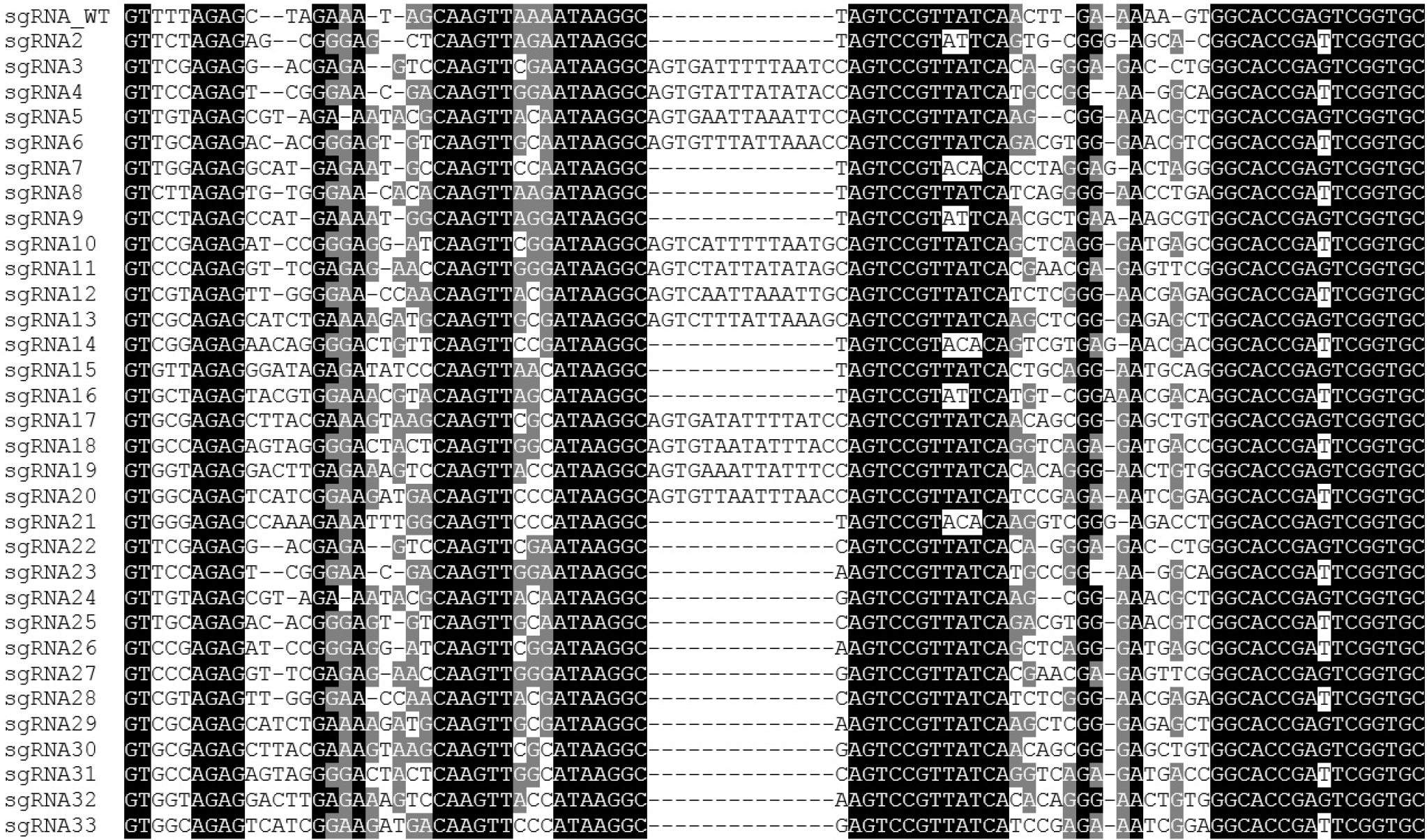
Complete list of sequence-divergent guide RNAs generated and assayed using the transcriptional activation reporter.

**Supplementary Figure 8.**
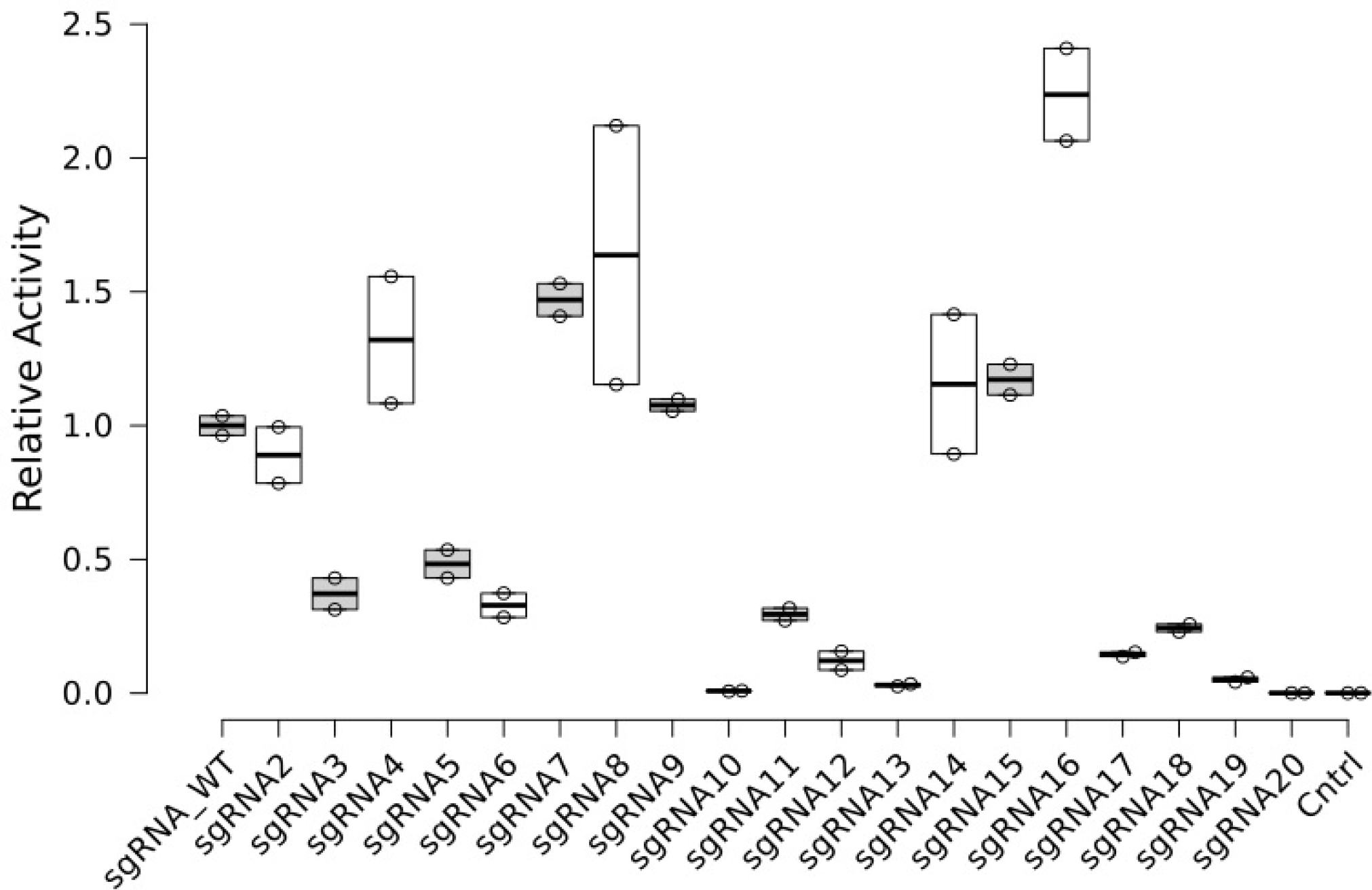
Results of the pilot screen of the first set of designed sgRNA sequences. #3-6, #10-13, and #17-20 all carried the extra insert; the latter 8 displayed markedly lower activity and were not further considered. The cause of the difference is unclear, although it is worth noting that these all had longer stem-loops than did #3-6, all of which were closer to the activity of the standard or ‘wild-type’ sgRNA.

**Supplementary Figure 9.**
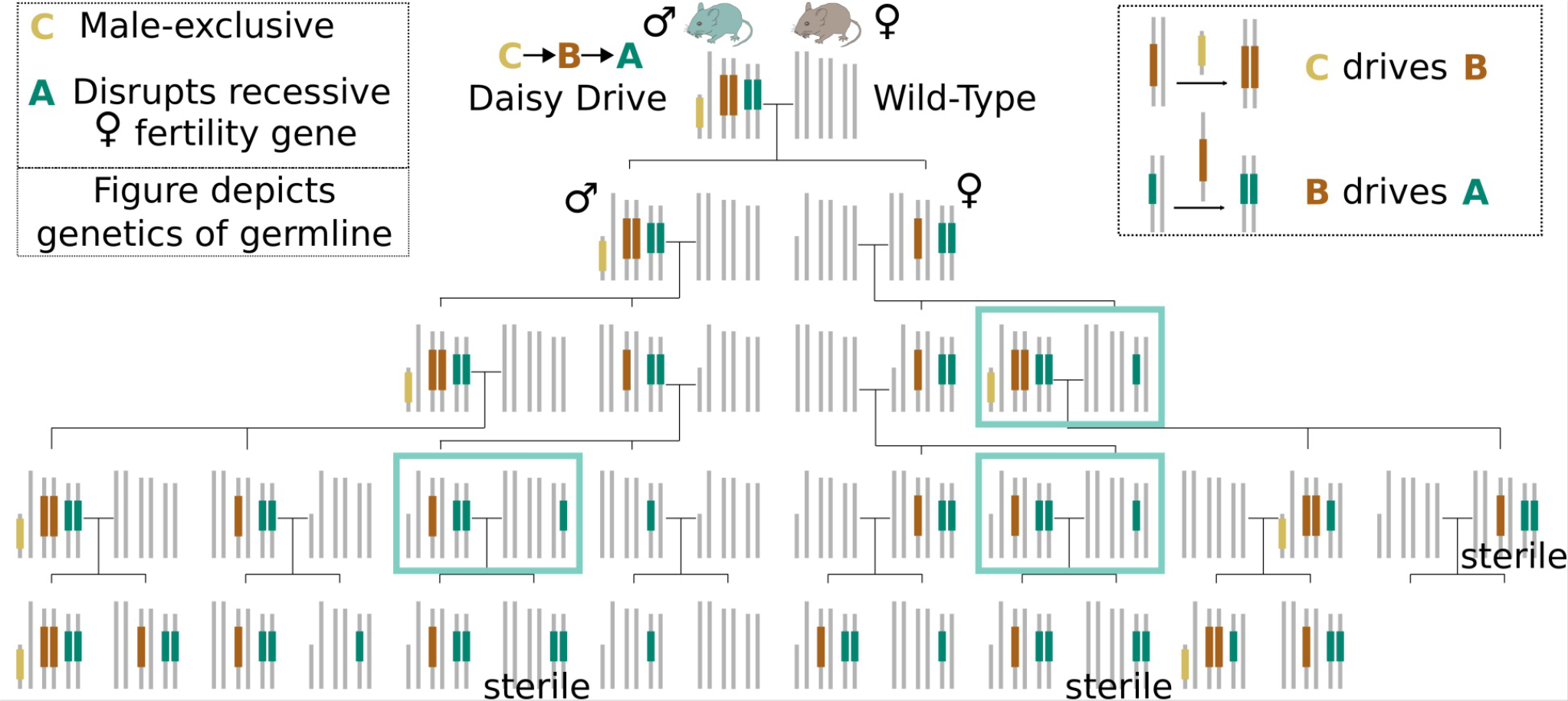
Potential family tree of a C→B→A genetic load daisy drive for which the payload in the A element disrupts a female fertility gene. The C element is male-linked, ensuring that it does not suffer a fitness cost from the loss of female fertility. Mating events between two parents carrying the A element (boxed) can produce sterile female offspring that will suppress the population. Males do not suffer a fitness cost due to disruption of female-specific fertility genes.

**Supplementary Figure 10.**
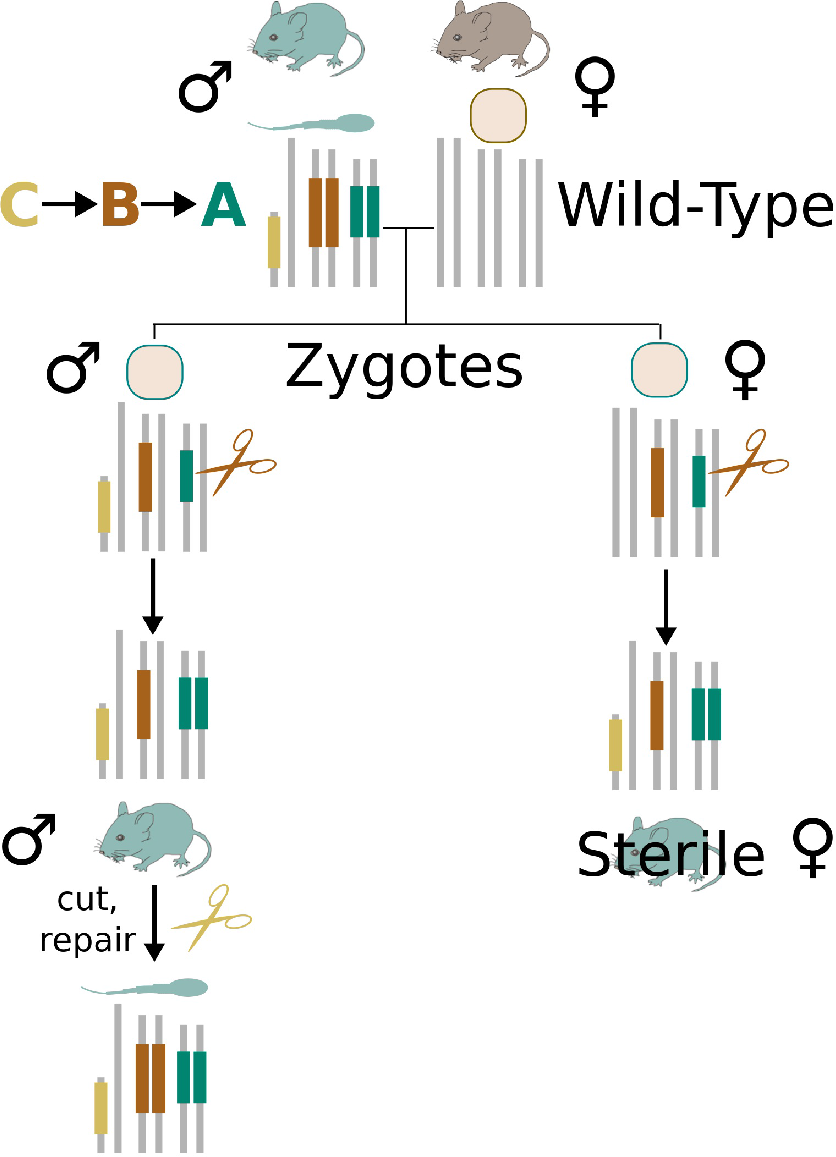
| A daisy-like system can create males with a male-linked sterile-daughter trait. Element (A) disrupts one or more recessive genes required for female fertility. The presence of a single copy of the B element causes the A element to drive in the zygote or early embryo, ensuring that females are sterile but leaving males unaffected. The C element is male-linked and causes the B element to drive, which can occur either in the germline as depicted or in the zygote or early embryo. The result is a male that produces sons like himself, but daughters that are sterile. These males should have fitness only slightly lesser than wild-type males and consequently should remain in the population for an extended period once released. Since population reproductive capacity depends directly on the number of sterile females, which in turn depends on the fraction of males with the sterile-daughter trait, this architecture could permit reversible titration of the local population by releasing sterile-daughter or wild-type organisms.

